# Identification of a set of endogenous retroviruses linked to IL-1-mediated immunopathology in severe COVID-19

**DOI:** 10.64898/2026.07.08.737320

**Authors:** Bessie Wang, Thomas Deckers, Eric Liu, Maria Tokuyama

**Affiliations:** Department of Microbiology and Immunology, Life Sciences Institute, The University of British Columbia, Vancouver, BC, Canada

## Abstract

SARS-CoV-2 infection triggers expression of endogenous retroviruses (ERVs), but whether this persists during COVID-19 and contributes to disease severity has not been well characterized. In this study, we analyzed bulk and single-cell RNA-sequencing datasets from multiple cohorts of severe COVID-19 patients to investigate the role of ERVs in COVID-19 immunopathology. We identified 33 differentially expressed proviral ERV loci in severe COVID-19 compared with healthy individuals (“severe COVID-19 signature ERVs”). ERV activation was associated with dysregulation of ERV epigenetic regulators, and their expression strongly correlated with key inflammatory pathways implicated in severe COVID-19, including neutrophil degranulation, interleukin signaling, and inflammasome activation. Anakinra treatment reversed activation of signature ERVs. Six signature ERVs were specific to intensive care unit (ICU) admission and significantly correlated with hospital-free days at day 45 (HFD-45). At the single-cell level, signature ERVs were significantly upregulated in erythroid-like and erythroid precursor cells and macrophages of patients with severe disease. Within these cells, we found evidence of ERV-associated differences in inflammatory gene expression, whereby cells expressing signature ERVs showed heightened expression of innate immune genes compared with cells not expressing these ERVs. Together, our study unmasked specific ERV loci activated in severe COVID-19 that are linked to immunopathology.

## Introduction

Severe acute respiratory syndrome coronavirus 2 (SARS-CoV-2) is a highly pathogenic human coronavirus that causes coronavirus disease 2019 (COVID-19)^1^. SARS-CoV-2 has impacted millions of people globally, and it continues to evolve and escape preexisting and vaccine-induced immunity, placing a sustained burden on healthcare systems. Early data from the pandemic indicated that while most infections resulted in mild to moderate symptoms, around 14 to 30 % of cases led to severe clinical outcomes, such as acute respiratory distress syndrome (ARDS), thrombosis, and multi-organ failure^2–4^. Although vaccination is highly effective in preventing severe disease, susceptible individuals develop complications that still require hospitalization^5^, highlighting the need to better understand factors that affect the severity of COVID-19.

The immunopathology of COVID-19 is characterized by a tipping of balance between sufficient viral control versus excessive inflammation and tissue damage^6^. Multiple factors are associated with severe outcomes, starting with the ineffective control of viral replication due to insufficient interferon (IFN) responses^7–10^, lower natural killer (NK) cell responses^11,12^, and maladapted cellular and humoral responses^13,14^. The impaired antiviral responses result in tissue damage, emergency myelopoiesis, and production of pro-inflammatory cytokines such as interleukin (IL)-6, IL-1β, IL-18, and TNF-α^15,16^. This in turn increases endothelial cell damage, vascular permeability, and neutrophil infiltration to pulmonary capillaries, which initiate a coagulation cascade that eventually causes coagulopathy and thrombosis that exacerbate COVID-19^17–19^.

Human endogenous retroviruses (HERVs) are a type of transposable element (TE) that make up approximately 8% of the human genome^20^ and are normally repressed through epigenetic silencing machinery^21,22^. However, ERVs are activated in healthy tissues and during disease, including infection by SARS-CoV-2 and other respiratory viruses^23,24^. In COVID-19 patients, ERVs are upregulated in the peripheral blood mononuclear cells (PBMCs) and bronchoalveolar lavage fluid (BALF)^25,26^. The presence of HERV-W, HERV-H, and HERV-K mRNA, as well as HERV-W envelope protein in COVID-19 patients are linked to disease severity^25–29^. However, the mechanisms and functional consequences of HERV activation in COVID-19 remain poorly understood.

Although HERVs are incapable of producing infectious viral particles in humans, there is mounting evidence supporting their immunomodulatory effects^30^. For instance, ERV long terminal repeats (LTRs) promote or enhance expression of proximal gene^31^, and ERV-derived long non-coding RNAs and enhancer RNAs modulate distal gene expression^32,33^. ERV envelope stimulates Toll-like receptor (TLR) 2 and 4^34^, while ERV-derived cytosolic RNA or DNA are sensed by retinoic acid-inducible gene I (RIG-I) or melanoma differentiation-associated protein 5 (MDA5) and cyclic GMP-AMP synthase (cGAS) to trigger an IFN response ^35–37^. This prompted us to ask whether ERV activation plays an immunomodulatory role that impacts disease severity in COVID-19.

We systematically analyzed the expression of proviral ERVs in the blood of severe COVID-19 patients to determine the specific ERVs that are activated in COVID-19, their cellular source, and links to immunopathology in COVID-19 using publicly available bulk and single-cell (sc) RNA-sequencing (RNA-seq) datasets. We identified key signature ERVs that are associated with severe COVID-19 (“severe COVID-19 signature ERVs”), which significantly correlated with key hallmarks of COVID-19 immunopathology, such as inflammasome activation and neutrophil degranulation. We also found elevated ERV expression, particularly in erythroid-like and erythroid precursor cells and macrophages, and this increase was accompanied by enhanced immune activation. Together, we provide evidence linking expression of a distinct set of ERVs to the immunopathology and disease severity of COVID-19.

## Results

### Locus-specific ERV analysis revealed a signature of ERVs defining severe COVID-19

To analyze ERV expression in severe COVID-19 patients, we used a publicly available whole blood RNA-seq dataset from a multicentric French COVID cohort^38^, consisting of healthy controls and patients classified as severe^39^. Using an updated version of ERVmap^40^, ERVmap2 (**Fig. S1A**), we identified 52 ERV loci that were significantly differentially expressed (DE-ERVs) in severe COVID-19 patients compared to healthy controls (**Fig. 1A**). We further filtered these DE-ERVs to remove potential signals coming from alternative transcriptional events, because many, though not all, of these DE-ERVs overlapped with genomic features such as introns, pseudogenes, and lncRNAs, or were near protein-coding genes (**Table S1**). These features could spill into ERVs by way of intron retention (IR) or transcriptional readthrough (RT) events, as evidenced by some of the DE-ERVs that extended beyond the proviral ERV region (**Fig. S1B**). A total of 19 ERVs that significantly correlated with overlapping genomic features and/or upstream genes were excluded (**Table S2**), resulting in a final set of 30 significantly upregulated ERV loci and 3 downregulated ERV loci in severe COVID-19 compared to healthy controls (**Fig. 1B and Table 1**). These ERVs were sufficient to segregate severe COVID-19 patients from healthy controls (**Fig. 1C**) and were henceforth designated “severe COVID-19 signature ERVs.”

**Figure 1.**
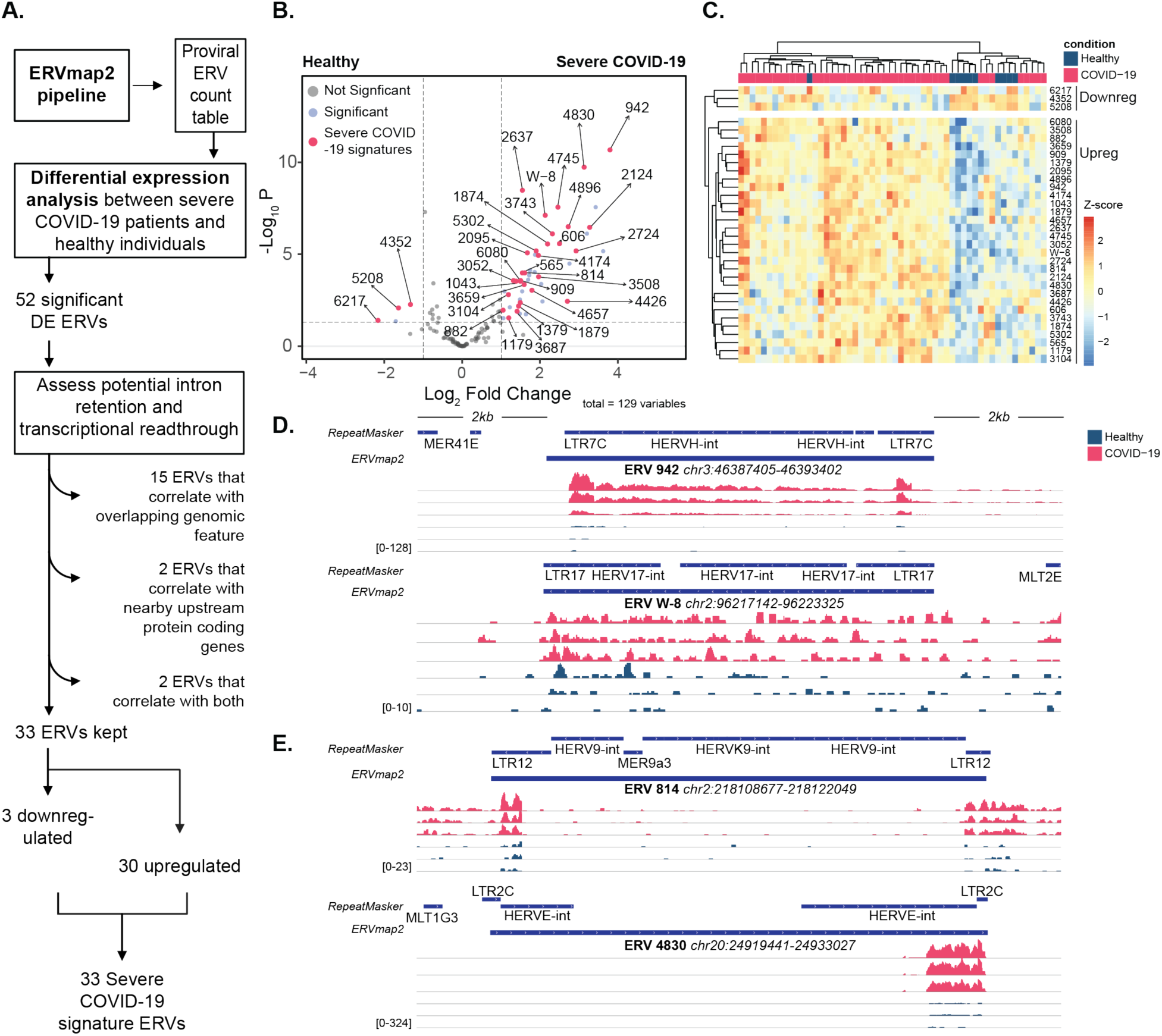
Identification of 33 differentially expressed proviral ERVs in severe COVID-19 patient blood. **(A)** Flow chart of workflow and filtering. **(B)** Volcano plot of differentially expressed ERVs. Blue, ERVs differentially expressed in COVID-19 patients (n=44) compared to healthy controls (n=10) (adjusted p < 0.05, |log2 FC| > 1.0); Pink, severe COVID-19 signature ERVs. **(C)** Heatmap of signature ERV expression in healthy and severe COVID-19 patients. **D, E** Read coverage of **(D)** ERV loci 942, W-8 and **(E)** ERV loci 814, 4830 in severe COVID-19 patients and healthy controls. Samples with the highest counts for each ERV locus in both conditions are displayed.

**Table 1.** Characteristics of 33 severe COVID-19 signature ERVs.

|  | erv_name | erv_group | supergroup | class | chromosome | start | end | strand |
| --- | --- | --- | --- | --- | --- | --- | --- | --- |
| Upregulated |  |  |  |  |  |  |  |  |
|  | 565 | HUERSP3 | HUERSP | 1 | 2 | 69789471 | 69799355 | - |
|  | 606 | THE | HSERVIII | 3 | 2 | 88876862 | 88890156 | + |
|  | 814 | HERV9 | HERVW9 | 1 | 2 | 218108677 | 218122049 | - |
|  | 882 | HERV9 | HERVW9 | 1 | 3 | 20269428 | 20281828 | + |
|  | 909 | HERVH | HERVHF | 1 | 3 | 32458603 | 32467714 | - |
|  | 942 | HERVH | HERVHF | 1 | 3 | 46387405 | 46393402 | - |
|  | 1043 | HERVT | MLLV | 1 | 3 | 101632044 | 101642053 | + |
|  | 1179 | HERVH | HERVHF | 1 | 3 | 152596603 | 152605968 | + |
|  | 1379 | HML6 | HML | 2 | 4 | 39539258 | 39544115 | - |
|  | 1874 | THE | HSERVIII | 3 | 5 | 70512459 | 70531584 | + |
|  | 1879 | HERV9 | HERVW9 | 1 | 5 | 72926261 | 72936085 | + |
|  | 2095 | HERVW | HERVW9 | 1 | 6 | 24667710 | 24680423 | - |
|  | 2124 | HERVIP | HERVIPADP | 1 | 6 | 35561856 | 35567823 | - |
|  | 2637 | HERVIP | HERVIPADP | 1 | 7 | 135174032 | 135184466 | + |
|  | 2724 | HERVH | HERVHF | 1 | 8 | 9058880 | 9065201 | - |
|  | 3052 | HERVH | HERVHF | 1 | 9 | 35019245 | 35025461 | - |
|  | 3104 | HML1 | HML | 2 | 9 | 90903903 | 90911679 | + |
|  | 3508 | HML3 | HML | 2 | 11 | 59923927 | 59931395 | + |
|  | 3659 | HERVH | HERVHF | 1 | 11 | 122824427 | 122834651 | + |
|  | 3687 | HERVE | HERVERI | 1 | 12 | 8281116 | 8291582 | - |
|  | 3743 | HERVH | HERVHF | 1 | 12 | 29135981 | 29140551 | - |
|  | 4174 | HML6 | HML | 2 | 14 | 69809151 | 69819704 | + |
|  | 4426 | HERVH | HERVHF | 1 | 17 | 11971743 | 11978102 | + |
|  | 4657 | HML5 | HML | 2 | 19 | 21620236 | 21631408 | + |
|  | 4745 | HERVH | HERVHF | 1 | 19 | 47392828 | 47398771 | + |
|  | 4830 | HERVE | HERVERI | 1 | 20 | 24919441 | 24933027 | + |
|  | 4896 | HERV3 | HERVERI | 1 | 21 | 38224030 | 38229830 | - |
|  | 5302 | HERVH | HERVHF | 1 | X | 12939864 | 12947330 | + |
|  | 6080 | HML8 | HML | 2 | 1 | 160687620 | 160690643 | - |
|  | W-8 | HERVW | HERVW9 | 1 | 2 | 96217142 | 96223325 | - |
| Downregulated |  |  |  |  |  |  |  |  |
|  | 4352 | HERVI | HERVERI | 1 | 16 | 20674232 | 20681995 | + |
|  | 5208 | HERV3 | HERVERI | 1 | Y | 21240221 | 21249982 | - |
|  | 6217 | HERVH | HERVHF | 1 | 1 | 229173970 | 229180173 | + |

Severe COVID-19 signature ERVs were 8.9kb in length on average and were distributed across 19 chromosomes (**Fig. S1C**). A large fraction of them were HERV-H, the most abundant ERV subfamily in the human genome^41^, but the expression of both evolutionarily older elements (HUERSP3, HML5, HML6, HERVIP, HERV9, and HERVE) and younger elements (HML3 and HERV-H) indicated that ERV expression in severe COVID-19 is independent of their age of integration. Contrary to previous reports, we did not observe significant upregulation of HERV-K (HML-2)^26,28^, despite our ability to detect coverage for most of the HML-2 loci using our pipeline (**Fig. S1D**).

We next assessed the read coverage of severe COVID-19 signature ERVs to better understand their transcriptional pattern. For several of the highly upregulated ERVs, such as 942 (HERV-H) and W-8 (HERV-W), reads covered the entire or nearly full length of the proviral sequence (**Fig. 1D**). In contrast, for ERVs such as 814 (HERV-9) and 4830 (HERV-E), read coverage was restricted to just the 5′ or the 3′ LTRs (**Fig. 1E**). In addition, based on predicted conserved protein domains of HERVs^42^, several severe COVID-19 signature ERVs contained highly conserved retroviral domains (**Table S3**), indicating potential for protein production from these loci. Overall, we observed distinct transcriptional characteristics of the severe COVID-19 signature ERVs.

### ERV expression in COVID-19 is associated with differential expression of epigenetic regulators

ERVs are epigenetically silenced by DNA methylation^43^ and trimethylated histone 3 lysine 9 (H3K9me3) deposition by tripartite motif-containing protein 28 (TRIM28)^44^ and the human silencing hub (HUSH) complex^45^. Ten-eleven translocation (TET) dioxygenases^46,47^ and peptidylarginine deiminases (PADIs)^48^ are also known to regulate ERV expression through removal of repressive marks. We hypothesized that the observed expression of ERVs is linked to changes in the expression of these epigenetic regulators.

We observed significantly lower expression of ERV repressors, including the HUSH complex components periphilin 1 (*PPHLN1*) and MORC family CW-type zinc finger 2 (*MORC2*) in severe COVID-19 patients compared to healthy controls (**Fig. 2A**). Components of the HUSH complex, transcription activation suppressor (*TASOR*) and methyl-H3K9-binding protein (*MPHOSPH8*), as well as *TRIM28* were not differentially expressed. The dioxygenases *TET2* and *TET3*, and the peptidylarginine deiminases *PADI2* and *PADI4* were significantly upregulated in COVID-19 compared to healthy. DNA methyltransferases *DNMT1* and *DNMT3A* showed no significant differences, while *DNMT3B* expression was significantly elevated in severe COVID-19 patients.

**Figure 2.**
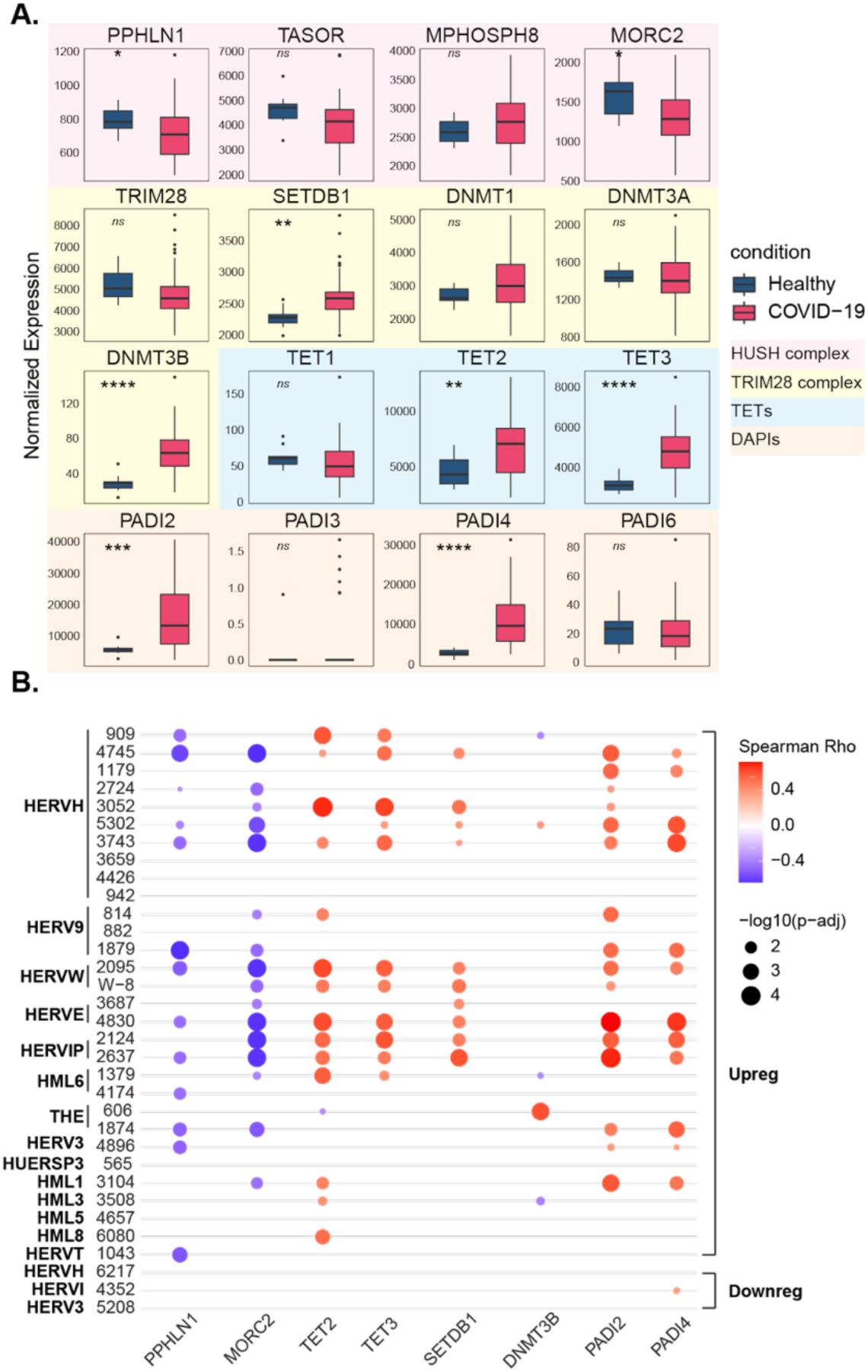
Severe COVID-19 signature ERVs correlate with epigenetic regulators in severe COVID-19 patients. **(A)** Expression of selected epigenetic regulators in severe COVID-19 patients compared with healthy individuals. Each box represents the interquartile range (IQR) of normalized gene expression, with the center line indicating the median. Whiskers extend to the most extreme data points within 1.5× the IQR; outliers beyond this range are shown as individual points. Statistical comparisons between conditions were performed using the Wilcoxon rank-sum test. Genes with significant expression differences are marked with asterisks: p < 0.05 (*), p < 0.01 (**), p < 0.001 (***), and p < 0.0001 (****). **(B)** Dot plot showing Spearman correlations between severe COVID-19 signature ERVs and epigenetic regulators. Dot color indicates the pairwise Spearman correlation coefficient, and dot size represents the adjusted p-value based on Benjamini–Hochberg FDR correction. Only significant correlations (adjusted p < 0.05) are displayed.

To better understand the link between these regulators and severe COVID-19 signature ERVs, we performed a Spearman correlation analysis. Of the 33 signature ERVs, 24 showed significant correlation (adjusted p-value < 0.05) with at least one of the differentially expressed regulators (**Fig. 2B**). Members of the HUSH complex, *PPHLN1* and *MORC2*, negatively correlated with signature ERVs, while *TET2*, *TET3*, *PADI2*, and *PADI4* positively correlated with ERV expression across multiple HERV families. Although the elevated expression of *SETDB1* and its positive correlation with ERVs were unexpected, our data suggest that the loss of silencing mediated by the HUSH complex, TETs, and PADIs is likely a key contributor to ERV expression in severe COVID-19.

### Severe COVID-19 signature ERVs correlate with inflammatory gene modules in COVID-19

ERVs modulate inflammatory responses through direct or indirect mechanisms^31,30^, thus we probed whether ERV expression directly correlates with inflammatory pathways in severe COVID-19 patients. Using weighted gene correlation network analysis (WGCNA)^49^, we identified 13 differentially expressed module eigengenes (MEs) in COVID-19 patients, of which, ME5, 48, 2, 61, 9, and 0 significantly correlated with the upregulated signature ERVs (**Fig. 3A**). We then performed a Reactome pathway enrichment analysis for these MEs and filtered the top five most enriched pathways for each ME (**Fig. 3B**). Next, we selected genes within the top five enriched Reactome pathways for each ME and found that Reactome genes from ME2 exhibited the strongest correlation with the signature ERVs (**Fig. 3C**). These included genes from the *Leishmania* infection pathway, interleukin (IL1, IL-8, IL7, IL2) signaling pathway, neutrophil degranulation pathway, as well as the TLR pathway. Notably, most of the Reactome genes in ME2 significantly correlated with ten or more of the signature ERVs (**Fig. S2**), suggesting that these ERVs are part of a regulatory network with inflammatory pathways that are involved in COVID-19.

**Figure 3.**
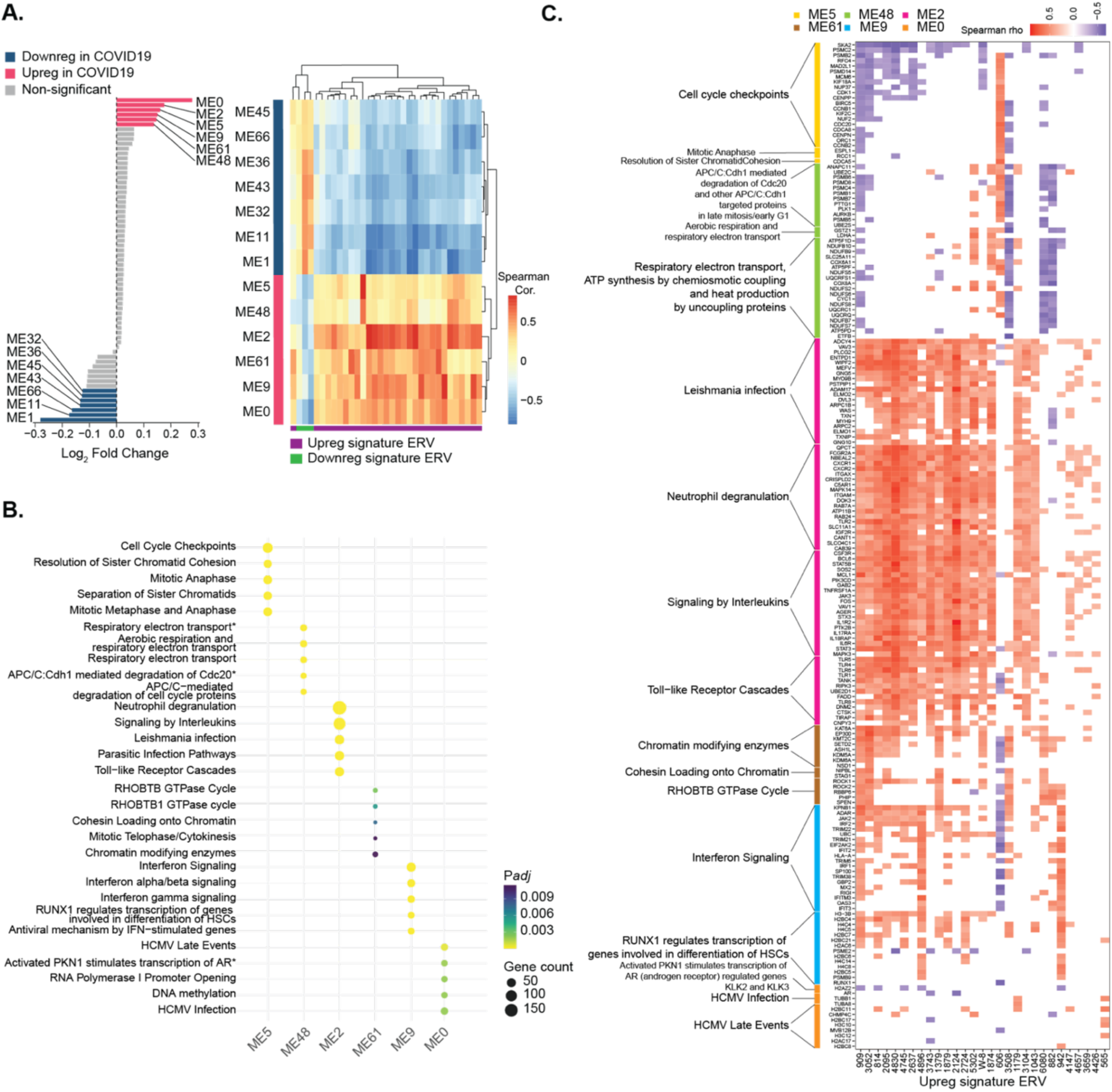
Severe COVID-19 signature ERVs are associated with inflammatory gene modules. **(A)** Left: Bar plot showing log₂ fold change of differentially expressed gene modules (FDR <0.05, |log2FC|>1) in severe COVID-19 compared with healthy controls. Right: Heatmap showing Spearman correlations between COVID-19–associated MEs and the signature ERVs; only significant correlations (adjusted p < 0.05) are shown. **(B)** Dot plot showing the top five enriched Reactome pathways for MEs upregulated in severe COVID-19. **(C)** Heatmap showing Spearman correlations between 30 upregulated signature ERVs and genes from the top five Reactome pathways. Duplicated genes across different reactome terms were assigned to the term with lowest adjusted p-value. The top 20 genes correlated with the largest number of ERVs are displayed, and only significant correlations (adjusted p < 0.05) are shown.

ME61 and 9 also showed positive correlation with upregulated ERVs, albeit with fewer ERVs (**Fig. 3C and Fig. S2**). Reactome genes from ME5, 48, and 0 mostly correlated with a single ERV and negatively correlated with signature ERVs (**Fig. 3C and Fig. S2**). Overall, we observed that ERV expression is linked to hallmarks of innate immune response to infection, which is known to contribute to the immunopathology of COVID-19.

### Increased ERV expression is associated with innate immune activation in severe COVID-19 and is responsive to anakinra treatment

Based on the expression of upregulated severe COVID-19 signature ERVs, COVID-19 patients segregated into two groups: an ERV-high group and an ERV-low group, the latter clustering more closely with healthy donors (**Figs. 1C and 4A**). We observed that ERV-high patients expressed higher levels of myeloid-dominant inflammatory genes involved in interferon signaling, neutrophil degranulation, parasitic infection pathway, RUNX1 signaling, interleukin signaling, and TLR cascade (**Fig. 4B**). Specifically, components of IL-1 and IL-18 signaling pathways correlated with a cluster of 23 upregulated signature ERVs, and these correlations were unique to the ERV-high group of patients (**Fig. S3A**). Moreover, in the ERV-high group, we found that ERVs strongly correlated with genes that are primarily involved in the priming stage (signal 1) of the NLRP3 inflammasome activation pathway (**Fig. S3B**), a major driver of COVID-19 pathogenesis^15,16,50^.

**Figure 4.**
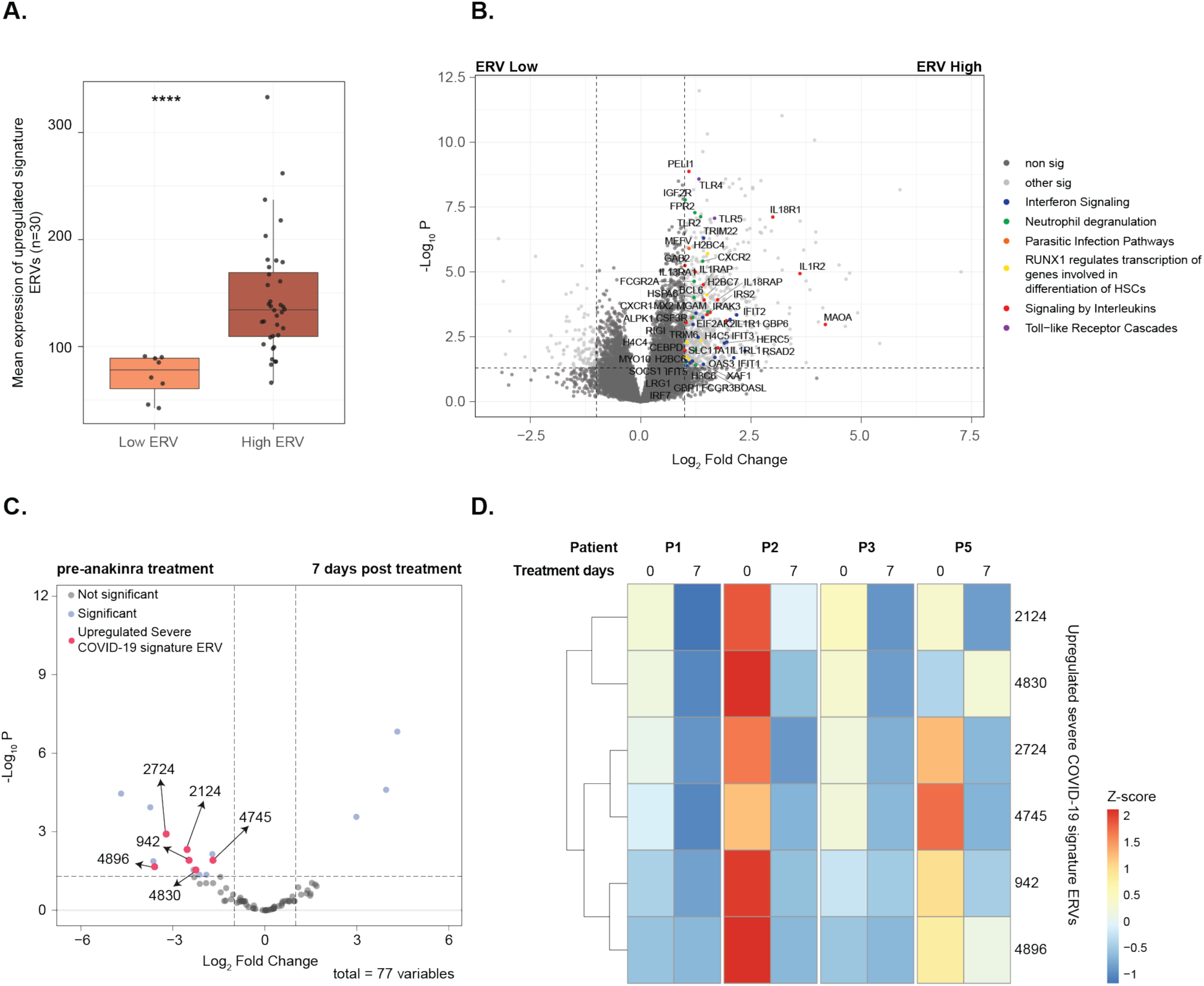
Severe COVID-19 patients with high ERV expression exhibit elevated innate immune gene expression. **(A)** Mean expression of the 30 upregulated signature ERVs in ERV-high (n=36) versus ERV-low (n=8) severe COVID-19 patients. Statistical comparisons between the two groups were performed using the Wilcoxon rank-sum test. Adjusted p < 0.0001 (****). **(B)** Volcano plot of differentially expressed (adjusted p < 0.05, |log2 FC| > 1.0) genes in ERV-high compared to ERV-low severe COVID-19 patients. Genes belonging to enriched Reactome pathways identified from gene modules upregulated in COVID-19 are highlighted. **(C)** Volcano plot of differentially expressed ERVs 7 days post anakinra treatment (n=4) in severe COVID-19 patients compared pre-treatment (n=4) (adjusted p < 0.05, |log2 FC| > 1.0); Pink, upregulated severe COVID-19 signature ERVs. **(D)** Heatmap of the expression of 6 upregulated signature ERVs pre and post treatment across patients.

To further define the link between ERVs and inflammasome activation, we analyzed ERV expression in severe COVID-19 patients before and after 7 days of treatment with the IL-1 receptor antagonist, anakinra^51^. Six upregulated signature ERVs were significantly downregulated following treatment (**Figs. 4D** and **4E**). The fold decrease in ERV expression was comparable to changes in clinical markers such as CRP and LDH in most patients (**Fig. S4**). Together, these findings provide evidence for a robust association between ERV activation and inflammasome activation in severe COVID-19.

### A subset of severe COVID-19 signature ERVs is associated with ICU admission status and disease severity

In order to test whether ERVs independently correlate with disease severity, we compared the expression of ERVs in COVID-19 patients, who were admitted to the intensive care unit (ICU) versus those who were not^52^. We identified fifteen DE-ERVs that were differentially expressed in ICU patients, and of these, eight DE-ERVs were defined as ICU signature ERVs upon filtering for overlaps and potential alternative transcriptional events, and six of them overlapped with the severe COVID-19 signature ERVs: ERV 2124, 3743, 1874, 5302, 4830, and 2724 (**Fig. 5A**).

**Figure 5.**
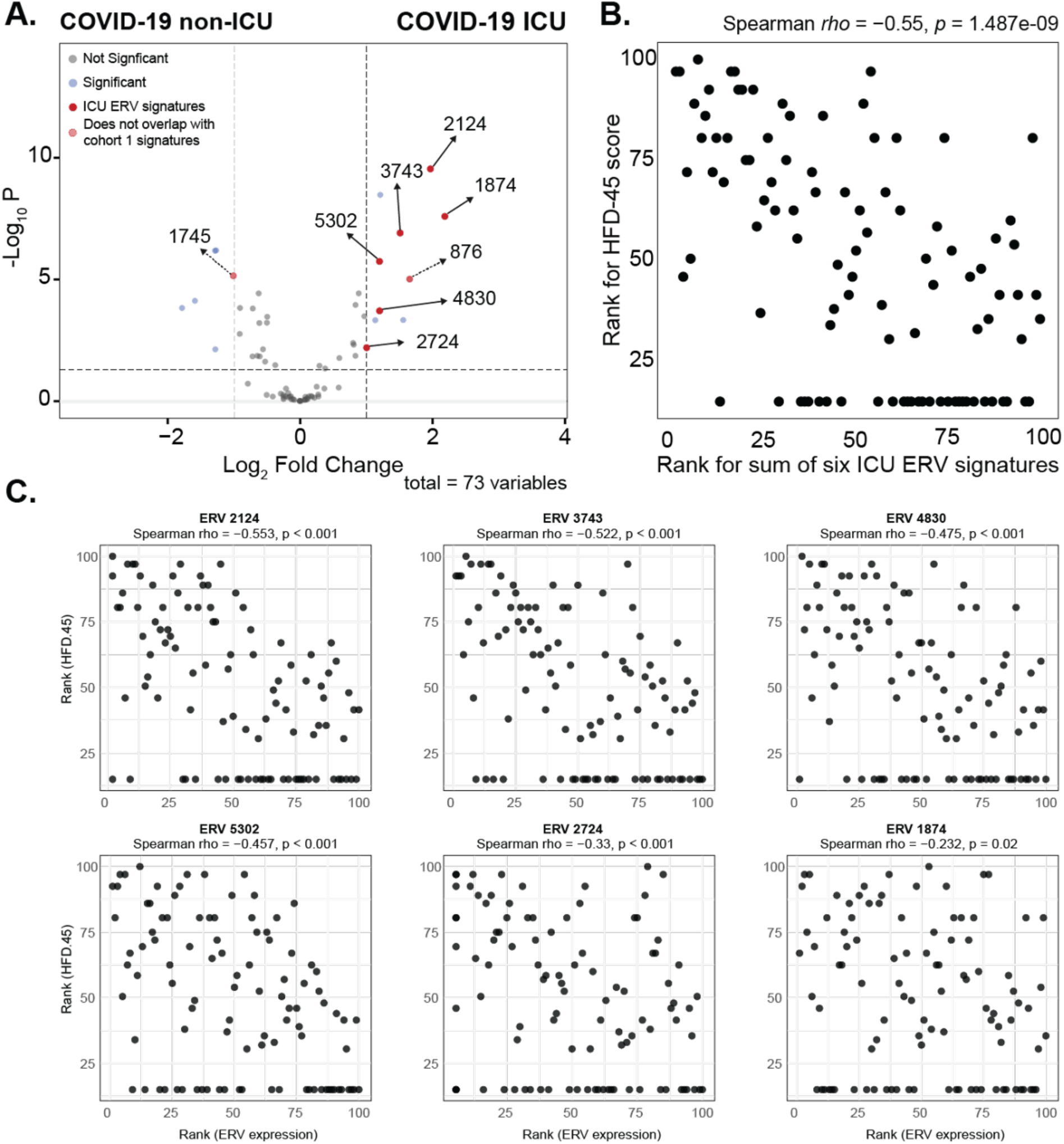
A subset of severe COVID-19 signature ERVs is specific to ICU status and correlates with disease severity. **(A)** Volcano plot showing differential expression of 73 ERVs between ICU (n=50) and non-ICU (n=50) COVID-19 patients. Blue, ERVs significantly elevated in ICU patients (padj < 0.05, |log2 FC| > 1.0); Red, ICU-specific signatures; Dotted line, ERV signatures that do not overlap with cohort 1. **(B)** Scatter plots of the ranked sum of the six ICU signature ERV expressions and HFD-45 score. **(C)** Scatter plots of ranked sum of individual ICU signature ERV expression and HFD-45 score.

To assess the relationship between these ICU signature ERVs and disease severity, we analyzed their association with hospital-free days at day 45 (HFD-45), a clinical metric of disease severity from the original study. The combined expression of the six ERVs negatively correlated with HFD-45 (r = −0.55, p < 0.05) (**Fig. 5B**), as did the individual ERVs (**Fig. 5C**). Notably, ERV 2124 had a stronger correlation with HFD-45 than the combined ICU ERVs. These results indicate that ICU signature ERVs are independent markers of poor clinical outcomes in COVID-19 patients, especially ERV 2124.

Lastly, to assess whether severe COVID-19 signature ERVs are specific to COVID-19, we compared ERV expression in severe COVID-19 patients to those who exhibited COVID-like symptoms but tested negative for SARS-CoV-2^52^. Ten ERVs were significantly upregulated in COVID-19 patients, whereby four were severe COVID-19 signature ERVs (565, 882, 2124, and W-8) and one was also an ICU signature ERV (2124) (**Figs. S5A and S5B**), revealing ERV loci associated with severe disease that are unique to COVID-19.

### Severe COVID-19 signature ERVs are expressed in erythroid-like and erythroid precursor cells and macrophages and correlate with innate immune activation

To further characterize the severe COVID-19 signature ERVs, we analyzed scRNA-seq data from the blood of severe COVID-19 patients^53^ using Cell Ranger and a customized genome reference file with stranded proviral ERV annotations (**Fig. 6A**). We identified ten distinct cell types in the blood of severe COVID-19 patients and healthy controls (**Fig. S6A**), and observed a decrease in T cells, NK cells, and macrophages and an increase in neutrophils, monocytes, and erythroid-like and erythroid precursor cells in COVID-19 **(Fig. 6B)**.

**Figure 6.**
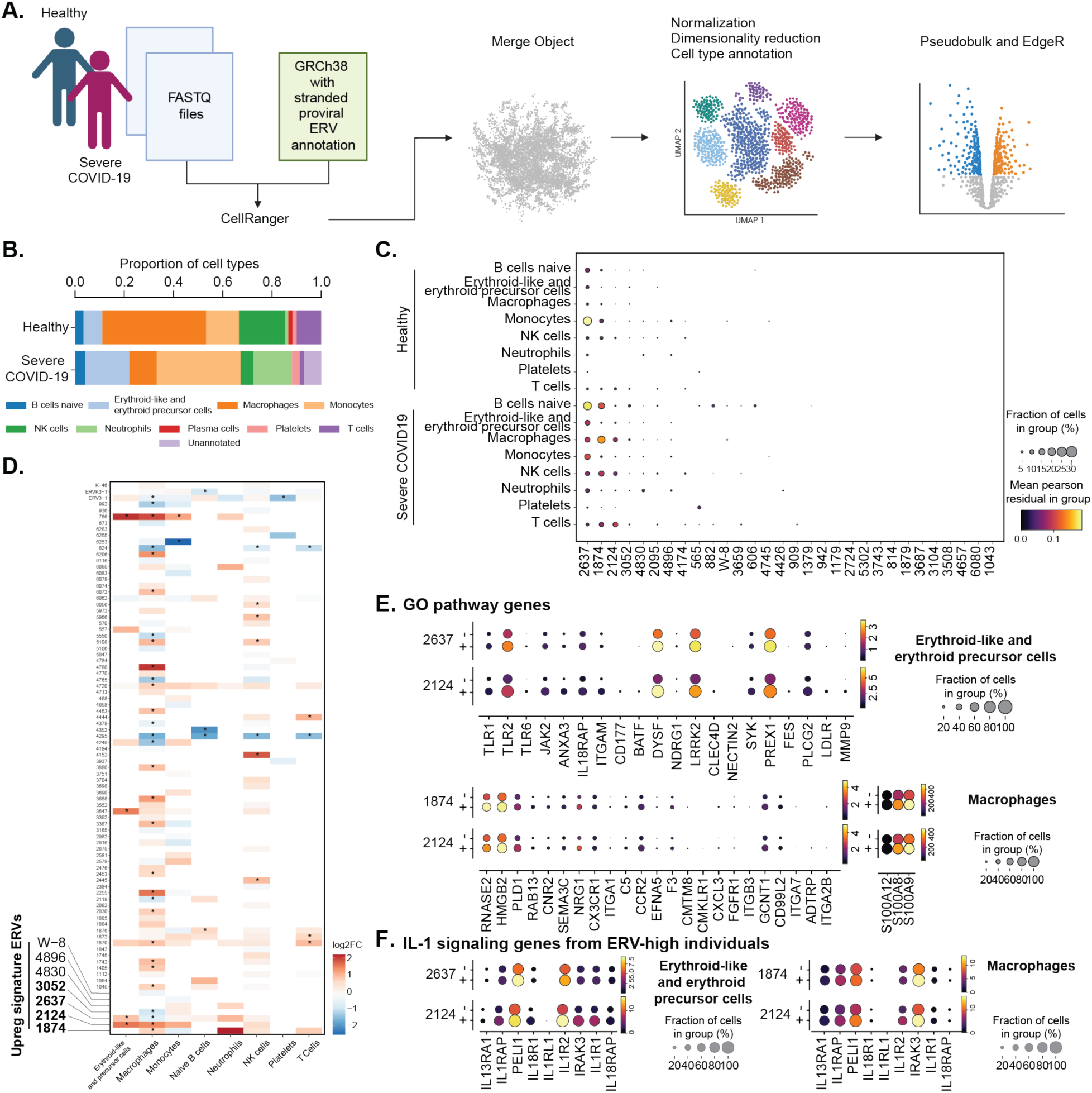
Expression of severe COVID-19 signature ERVs is strongly linked to innate immune activation at the single-cell level. **(A)** Workflow of scRNAseq analysis. Created with BioRender.com. **(B)** Bar plot showing the proportions of identified cell types in healthy controls (n = 6) and severe COVID-19 patients (n = 20). **(C)** Dot plot showing the expression levels and within-group proportions of cells expressing the 30 upregulated severe COVID-19 signature ERVs across cell types in healthy controls and severe COVID-19 patients. **(D)** Heatmap showing the Log2 fold change of differentially expressed ERVs in severe COVID-19 compared to healthy from the pseudobulk analysis. Asterisks indicate significant difference (adjusted p < 0.05). Dot plot comparing the expression and within-group proportions of cells expressing **(E)** GO pathway genes and (**F**) IL-1 signaling genes in erythroid-like and erythroid precursor cells and macrophages that either express (+) or do not express (–) signature ERVs.

We next assessed the expression of the upregulated severe COVID-19 signature ERVs in each cell type, except for the unannotated cluster and the plasma cell cluster that were too low in abundance. While 13 out of the 30 upregulated severe COVID-19 signature ERVs fell below the threshold of detection (fewer than 1% of cells) in either healthy or COVID-19, the remaining 17 signature ERVs were detected in up to 21% of cells in severe COVID-19 patients (**Fig. 6C**). Among these, ERV 2637, 1874, and 2124 were detected in all cell types, while other ERVs exhibited cell-type-specific expression patterns: ERV 565 in platelets, 882 and 606 in naive B cells, 4426 in neutrophils, and 909 in T cells.

To quantify differential expression of ERVs in these cell types between severe COVID-19 patients and healthy controls, we performed a pseudobulk analysis followed by a differential expression analysis within each cell type. In total, 88 ERVs were detected in severe COVID-19 patients, most of which were differentially expressed in at least one cell type (**Fig. 6D**) and included 4 severe COVID-19 signature ERVs that were significantly differentially expressed in severe COVID-19 compared to healthy controls. ERV 2637 and 2124 were upregulated in erythroid-like and erythroid precursor cells, ERV 2124 and 1874 were upregulated in macrophages, and ERV 3052 and 2637 were downregulated in macrophages.

We next probed whether heightened expression of ERV 2637, 2124, and 1874 in erythroid-like and erythroid precursor cells and macrophages is associated with the inflammatory state of these cells. Gene ontology (GO) enrichment analysis of differentially expressed genes revealed enrichment of immune-related pathways in these cells (**Fig. S6B**). We then segregated cells based on their expression of ERV 2637, 2124, or 1874 and compared the expression of these immune-related genes in cells that expressed (+) or did not express (-) ERVs in severe COVID-19 patients. In erythroid-like and erythroid precursor cells, cells expressing ERV 2124 and 2637 expressed higher levels of *TLR2*, *DYSF*, *LRRK2*, and *PREX1* in greater proportion of cells. In addition, a higher proportion of ERV+ erythroid precursor cells expressed *TLR1*, *JAK2*, *ANXA3*, *IL18RAP*, *SYK*, and *PLCG2* compared to ERV-cells (**Fig. 6E**). Similarly, macrophages expressing ERV 2124 and 1874 exhibited higher expression of chemotaxis-related genes compared to ERV-cells. We further examined the expression of epigenetic regulators of ERVs in these cell types and found that ERV+ cells showed increased expression of *TET2*, *TET3*, *PADI2*, and *PADI4* compared to ERV-cells, in both cell types (**Fig. S6C**).

Finally, we examined whether the associations between ERV expression and IL-1 signaling pathway genes or NLRP3 inflammasome-associated genes identified in our bulk RNA-seq analysis were also present in erythroid-like cells and erythroid precursor cells and macrophages. ERV+ cells showed increased expression of IL-1 signaling genes (**Fig. 6F**) and NLRP3 inflammasome-associated genes compared with ERV-cells (**Fig. S6D**). Together, these data demonstrate a clear link between ERV expression and the immunopathology of COVID-19 at the single-cell level.

## Discussion

ERVs are transcriptionally active in human tissues and are dynamically regulated under stress and disease conditions^23,30,54^. Despite significant improvements in bioinformatic pipelines, these workflows remain imperfect for the purpose of quantifying locus-specific proviral ERV expression independently of other genomic features. Here, we updated the ERVmap pipeline ^40^ and incorporated steps to reduce signals arising from likely alternative transcriptional events.

Specifically, ERVs showing strong correlation with nearby or overlapping genomic features were excluded from downstream analysis. Although the use of a less stringent threshold or other methods that specifically test for IR and RT could have yielded more DE-ERVs, these modifications increased the signal-to-noise ratio for quantification of locus-specific proviral ERV expression in bulk short-read RNA-seq data.

Several bioinformatic tools can quantify TEs in scRNA-seq data^55–57^, but they have not been widely applied for detection of proviral ERVs in a locus-specific manner. Using a customized Cell Ranger workflow together with scTransform normalization^58^ and pseudobulk analyses^59^, we detected cell-type-specific ERV expression despite the limited read depth of scRNA-seq. However, many signature ERVs remained below the limit of detection, which is an inherent limitation of single-cell approaches. Our study shows that pairing bulk RNA-seq outputs with scRNA-seq analysis provides a robust framework for identifying disease-associated ERVs.

COVID-19 triggers global changes in DNA methylation and histone modifications, which are strongly associated with disease severity^60,61^. Consistent with previous studies^62,63^, *TET2/3* and *PADI2/4* were significantly elevated in severe COVID-19 patients, and these factors correlated with the expression of severe COVID-19 signature ERVs. These findings are consistent with established roles of TET enzymes in DNA demethylation^64^ and ERV regulation^46,47^, as well as the ability of PADI4 promote proviral transcription of HIV^65^. However, ERV expression negatively correlated with components of the HUSH complex, and we did not observe associations with DNMTs or TRIM28. In addition, we speculate that the immunoregulatory effects of SETDB1^66^ possibly overshadowed its effect on ERVs in this context. Nonetheless, our data suggest that epigenetic remodeling during COVID-19 likely enables the activation of ERVs.

Previous studies have reported activation of ERVs during acute SARS-CoV-2 infection or upon *in vitro* infection^67,68^. We observed activation of ERVs in COVID-19 patients sampled approximately 3 days after hospitalization or 7 to 14 days after symptom onset, suggesting that ERV expression persists beyond the acute phase of infection and likely does not require ongoing viral replication. Whether this reflects persistent inflammation and epigenetic remodeling or is a cause of the immune dysregulation, or both, remains to be addressed.

Severe COVID-19 signature ERVs correlated with genes associated with neutrophil degranulation and interleukin signaling, implicating them in the inflammatory and tissue damage responses that exacerbate COVID-19^15,19,69^. The link between ERV expression and inflammasome activation suggests either a co-regulated response or a potential causal relationship, which was further supported by data showing downregulation of signature ERVs following anakinra treatment. Together with our finding that a number of these signatures correlate with ICU admission status and disease severity metric, the ERVs identified in this study may serve as molecular biomarkers of disease severity. Moreover, similar links between ERVs and inflammasome activation may be observed in other post-acute infection sequelae.

Finally, our scRNA-seq analyses revealed erythroid-like and erythroid precursor cells, macrophages, and neutrophils as key cellular sources of several severe COVID-19 signature ERVs. Cells expressing these ERVs exhibited higher expression of innate immune response genes compared with ERV-negative cells within the same cell type, providing, to our knowledge, the first evidence linking ERV expression with immune activation at the single-cell level. These findings suggest that ERVs may participate in the expansion of erythroid precursor cells upon infection^70,71^ or affect local and systemic macrophage responses^72^. Future studies will be needed to directly probe the links between ERVs and immune activation within these cell populations.

Despite providing some new insights into the involvement of ERVs in COVID-19 immunopathogenesis, this study has several limitations. First, although we sought to match demographic and clinical severity features across the three cohorts, samples were not fully matched for potential confounding variables, including age and sex. The datasets also lacked information on whether patients had active viral replication at the time of sampling. Second, because the alternative transcription filtering applied to bulk RNA-seq data could not be directly implemented in the scRNA-seq analysis, we focused only on bulk-defined signature ERVs, leaving other potentially relevant ERV loci unexplored. Finally, our findings suggest that ERV activation may be associated with an exacerbated inflammatory state, further studies are needed to elucidate the mechanisms by which ERVs contribute to immune activation.

Vaccines have significantly reduced the burden of COVID-19 and associated complications. However, as the virus continues to evolve and attempts to escape preexisting immunity, understanding factors that drive severe outcomes remains important. Our study provides evidence linking specific ERVs to COVID-19 immunopathology and offers a framework for further investigations into their roles in COVID-19 as well as other inflammatory conditions.

## Methods

### Data collection

Publicly available bulk and single-cell RNA-seq datasets generated from whole blood samples of severe COVID-19 patients and healthy controls were obtained from the Gene Expression Omnibus (GEO) database. Bulk RNA-seq raw data for three cohorts were downloaded from accession numbers GSE171110^38^, GSE163317^51^, and GSE157103^52^. Single-cell RNA-seq raw data were downloaded from GSE157344^53^. All cohorts included both male and female participants and were comparable with respect to time from illness onset, median age, and disease severity criteria. We specifically selected datasets collected in 2020 or datasets whose metadata indicated that participants were unvaccinated to ensure the acquisition of pre-vaccination SARS-CoV-2 infection samples.

### Analysis of Bulk RNA-seq data

Raw RNA-seq data were processed using the ERVmap2 pipeline (https://github.com/Bessie92/ERVmap2) to obtain ERV and gene count matrices across all samples. A custom STAR genome index was generated by incorporating 3217 stranded proviral ERV annotations from ERVmap2 into the GRCh38.p14 reference genome as exon features. ERV features that overlapped between ERVmap2 annotation and the default genome annotation were removed from the genome annotation but kept in the ERVmap annotation, including ERV3-1, ERVFRD-1, ERVK3-1, ERVV-1, and ERVV-2. Individual transcripts with more than 90% overlap with ERVmap2 annotations were also excluded from the genome annotation file, including ERVW-1-202, ERVH48-1-201, ERVW-1-204, and ACSM3-215.

Sequencing reads were then aligned to the custom genome using STAR v2.7.9^73^ with the following parameters --outSAMtype BAM SortedByCoordinate --quantMode GeneCounts -- outFilterMultimapNmax 1 --outSAMmultNmax 1 --outSJfilterReads Unique -- outSAMstrandField intronMotif --outFilterMismatchNoverReadLmax 0.01 --outSAMattributes All XS. Reads mapping to multiple genomic loci were excluded. Resulting BAM files were sorted and indexed using samtools^74^. Counts for cellular genes and ERVs were generated using STAR’s --quantMode GeneCounts option and separated into distinct count files.

Raw cellular gene counts were used to estimate sample-specific size factors with DESeq2^75^. These size factors were subsequently applied to normalize both cellular gene and ERV counts. Normalized counts were used for downstream analyses and visualizations, including heatmaps and correlation analyses.

Differential expression analyses of ERVs and genes were performed using unnormalized counts using DESeq2^75^. Features with a *baseMean* value below 10 were removed from post-DESeq2 analysis. ERVs and genes were considered significantly differentially expressed if they had an absolute log2 fold change greater than 1 and an adjusted p-value below 0.05.

### Defining signature ERVs

The reference genome was intersected with ERV annotations using a custom script (erv_gtf_intersect.sh) to identify genomic features overlapping ERV loci and the nearest upstream protein-coding gene within 4 kb. Spearman correlations were calculated between DE-ERVs and each overlapping genomic feature. For ERVs overlapping multiple features on either strand, each ERV-feature pair was analyzed separately. ERVs showing strong correlations (adjusted p-value < 0.05, Spearman’s rank correlation coefficient (ρ) > 0.5) were excluded. The same correlation analysis was applied to DE-ERVs with nearby upstream protein-coding genes, and ERVs with significant correlations were filtered out.

### Protein domain prediction

A retroviral-like domain prediction database (HERV_domains_v5.bed) from HERVarium^42^ was used to intersect with the genomic coordinates of the 33 severe COVID-19 signature ERVs using BEDTools^76^ (*bedtools intersect -a sig_ervs.bed -b HERV_domains_v5.bed -s -wa -wb*).

### WGCNA

A gene co-expression network was constructed using the WGCNA R package^49^. Outlier genes were identified from the normalized gene expression matrix using the *goodSampleGenes* function. The matrix was then transformed into a weighted adjacency matrix and a corresponding topological overlap matrix (TOM) using the Pearson correlation method with a soft power (β) of 3. Average linkage hierarchical clustering was performed based on TOM dissimilarity (1–TOM), and gene modules were identified using a dynamic hybrid tree-cutting algorithm. These steps were implemented using the *blockwiseModules* function with the following parameters: *power = 3, networkType = “signed”, TOMType = “signed”, minModuleSize = 30,* and *maxBlockSize = 8000.* Module eigengenes (MEs) were tested for association with disease status using a limma linear model^77^, followed by empirical Bayes moderation (eBayes) to stabilize variance estimates across modules (adjusted p-value < 0.05 were considered significantly associated with COVID-19).

### Analysis of single-cell RNA-seq data

Single-cell RNA-seq raw data were processed using Cell Ranger (v8.0.1). The GRCh38.p14 genome annotation used for the ERVmap2 pipeline with proviral ERVs added as exon features, was filtered with the CellRanger *mkgtf* function to remove pseudogenes and used to generate a custom genome reference. Read alignment, UMI counting, and droplet filtering were performed with Cell Ranger using default parameters.

Downstream data analysis and visualization was performed using scanpy (version 1.11.4)^78^. Quality control excluded cells expressing fewer than 200 genes and cells with more than 13% mitochondrial gene content. Raw UMI counts for cellular genes were normalized by applying regularized negative binomial regression using SCTransform^58^. Pearson residuals were used to identify the top 3,000 highly variable genes, which were then used for dimensionality reduction by principal component analysis (PCA). A nearest-neighbor graph was constructed using the first 50 principal components and 50 nearest neighbors, followed by UMAP visualization and Leiden clustering. To determine an appropriate clustering resolution, Leiden clustering was repeated using resolution parameters of 0.02, 0.10, 0.30, 0.50, 0.80, and 1.50. Cluster assignments were visualized on the UMAP and compared across resolutions, and a resolution of 0.50 was selected for downstream analysis.

Cell types were annotated using the decoupler package^79^ calculating cell type enrichment scores using the ulm method (*dc.mt.ulm*) with human blood and immune system marker genes from PanglaoDB^80^.

One pseudobulk sample was generated per patient per cell type by aggregating cells from the corresponding subpopulation and calculating the mean gene expression within that subpopulation. Samples < 30 cells in a given cell type were excluded from the analysis for that population. Differential expression analysis was performed using edgeR^81^ on pseudobulk count matrices. Genes with low counts were removed using *filterByExpr*, followed by library-size normalization using TMM normalization with *calcNormFactors*. Dispersion was estimated with *estimateDisp*, and gene-wise negative binomial generalized linear models were fitted using the quasi-likelihood framework with *glmQLFit*.

### Statistical analysis

The statistical tests applied to each comparison are specified in the corresponding figure legends. All analyses were carried out in R, with statistical significance determined using a Benjamini– Hochberg adjusted p-value threshold of < 0.05.

## Code availability

Code used to perform the analyses and generate the figures presented in this study is available at https://github.com/Bessie92/Wang-Tokuyama_ERVs-in-severe-COVID-19. For pre-publication access, please contact the corresponding author.

## Author Contributions

Conceptualization: BW, MT

Methodology: BW, TD, and EL

Formal analysis: BW

Writing - original draft: BW

Writing – review & editing: BW, MT

Visualization: BW

Resources: MT Supervision: MT

## Acknowledgements

We acknowledge that this work was performed on the UBC Point Grey (Vancouver) campus, which sits on the traditional, ancestral, unceded territory of the xʷməθkʷəy̓əm (Musqueam) First Nation. This research was supported in part through computational resources and services provided by Advanced Research Computing at the University of British Columbia. We thank the IT support staff at UBC Department of Microbiology and Immunology for their help.

