## Supplementary Figures for "Identification of a set of endogenous retroviruses linked to IL-1-mediated immunopathology in severe COVID-19"

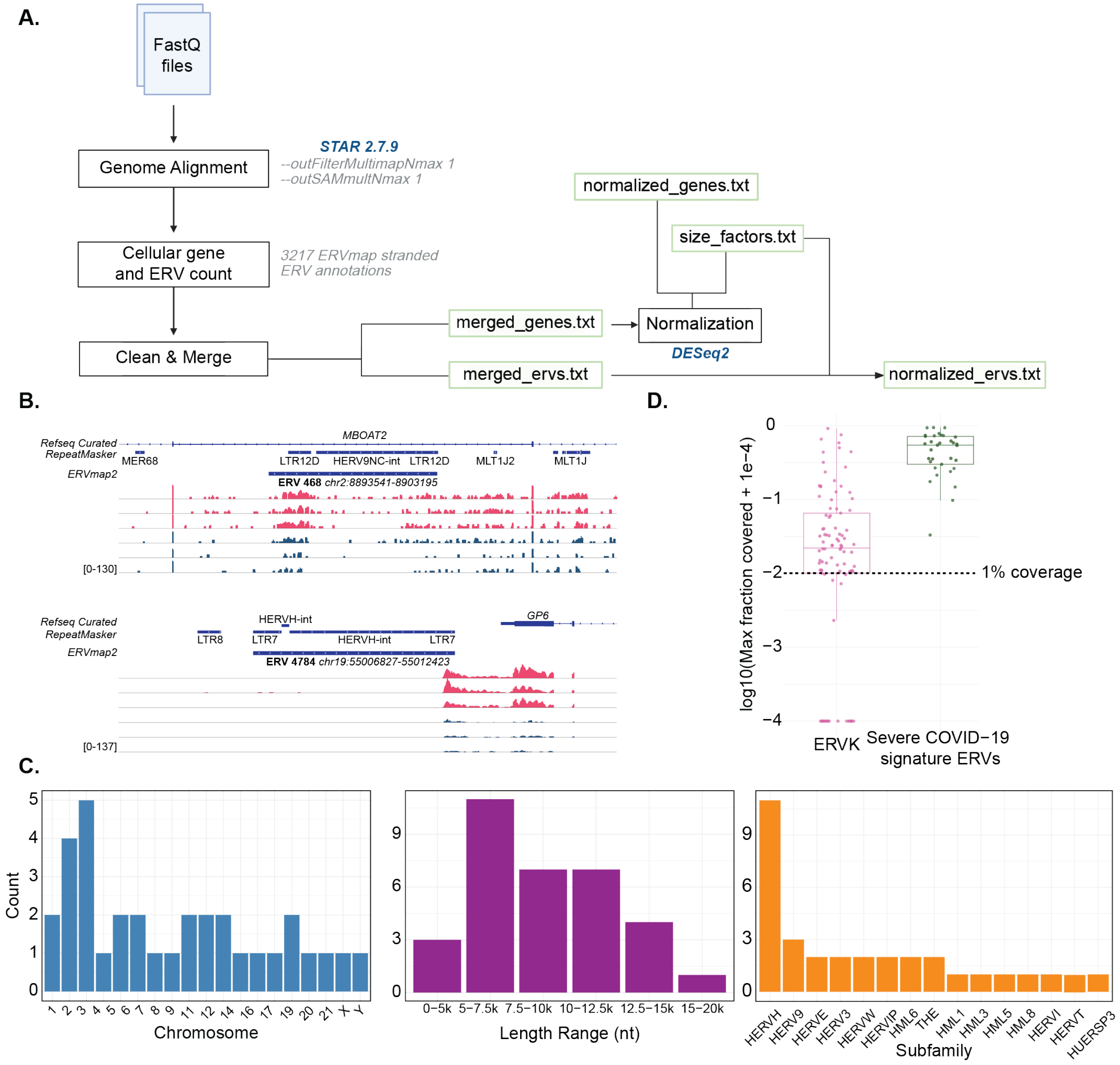


**Fig. S1. Characterization of ERVs detected in severe COVID-19 patients. (A)** Workflow of the ERVmap2 pipeline. **(B)** Read coverage for ERV 468 and 4784, illustrating non-ERV-specific expression patterns in severe COVID-19 patients (pink) and healthy controls (dark blue). **(C)** From left to right: histograms showing the chromosomal distribution, length, and group classification of the 33 severe COVID-19 signature ERVs. **(D)** Comparison of maximum fraction coverage between ERVK loci (n = 93, pink) and severe COVID-19 signature ERVs (n = 33, green) across all samples in the cohort.

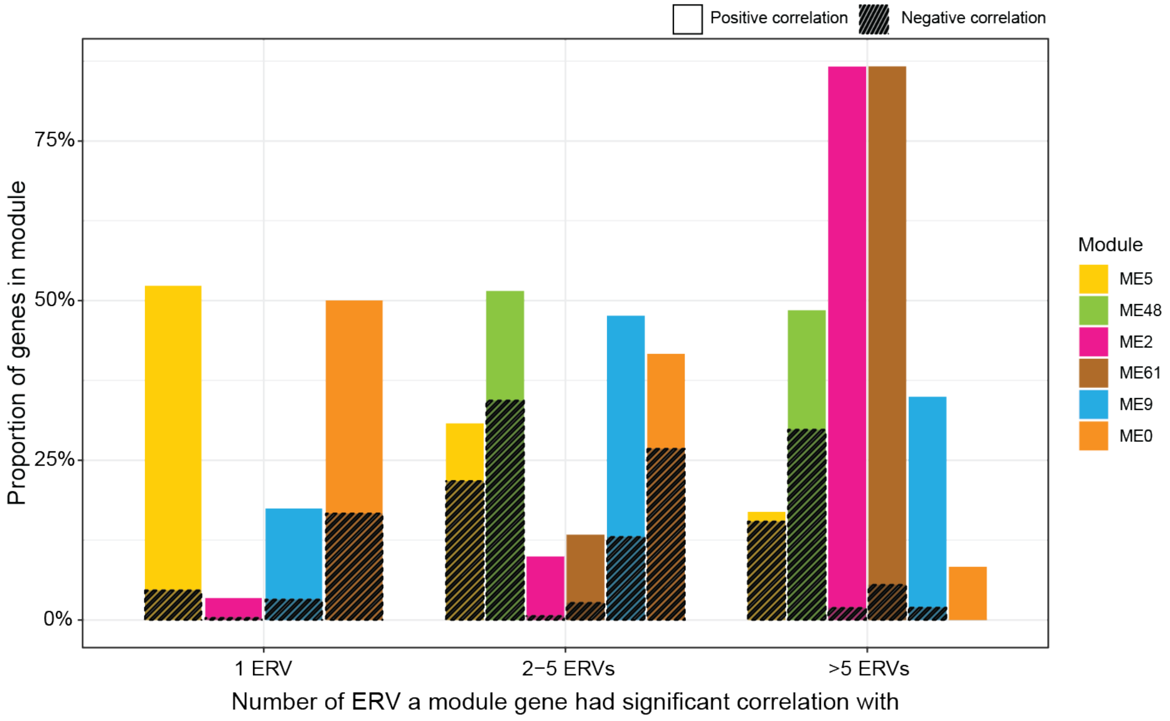


**Fig. S2. ME2 genes correlate with the largest number of signature ERVs among all module eigengenes.** Bar plot showing the proportion of genes representing the top five enriched Reactome pathways for each ME that significantly correlated (positively or negatively) (adjusted $p < 0.05$) with upregulated signature ERVs in severe COVID-19.

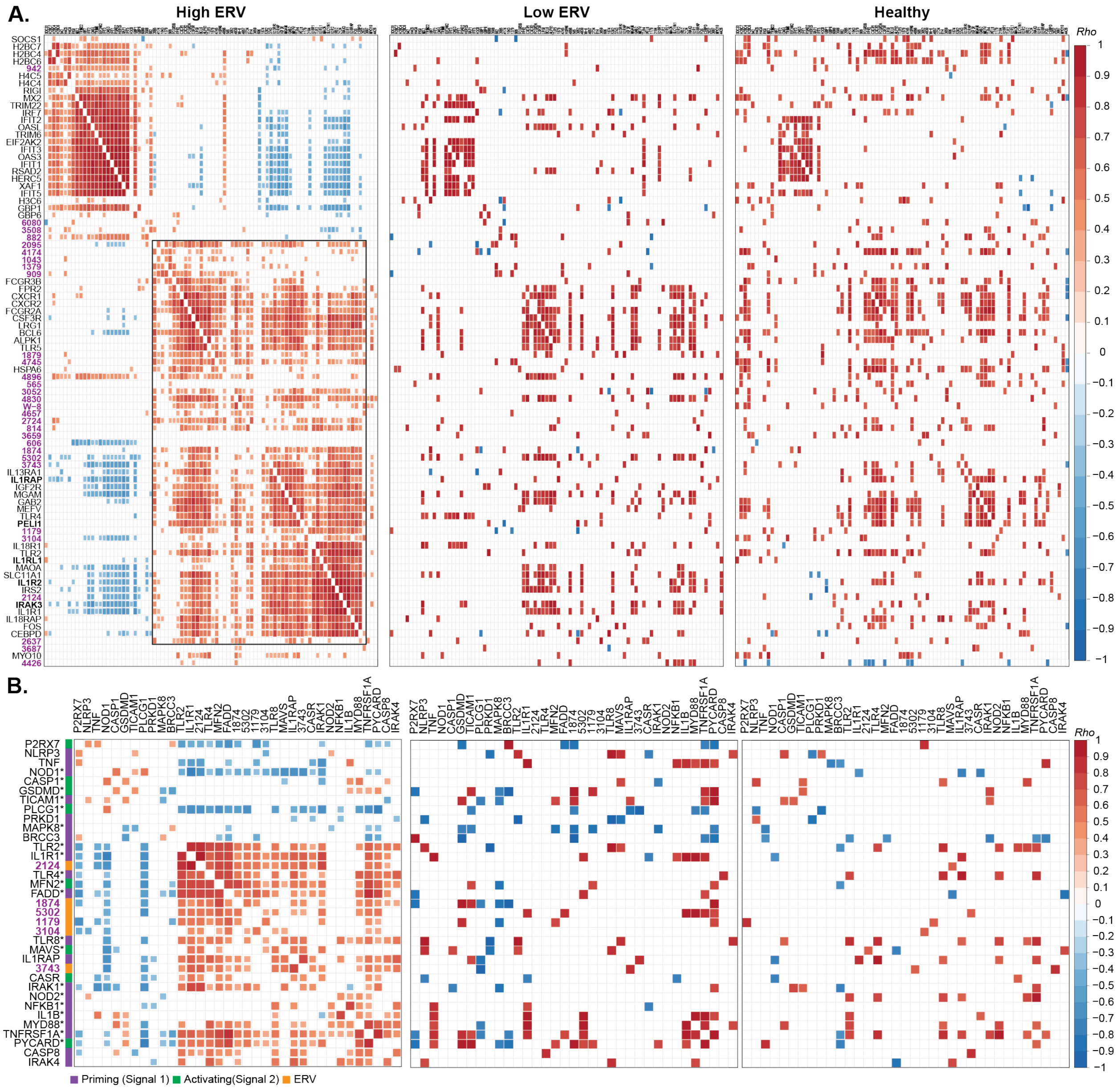


**Fig. S3 Upregulated signature ERVs correlate with inflammatory genes in severe COVID-19 patients. (A)** Heatmap showing Spearman correlations between upregulated Reactome genes and the 30 upregulated signature ERVs in ERV-high, ERV-low, and healthy groups. ERV names are shown in purple, and IL-1 signaling genes are highlighted in bold. **(B)** Heatmap showing Spearman correlations between ERVs and NLRP3 inflammasome activation genes in severe COVID-19 patients, grouped by ERV-high, ERV-low, and healthy groups. Genes upregulated in severe COVID-19 compared with healthy controls were annotated with asterisks (Wilcoxon rank-sum test, $p < 0.05$).

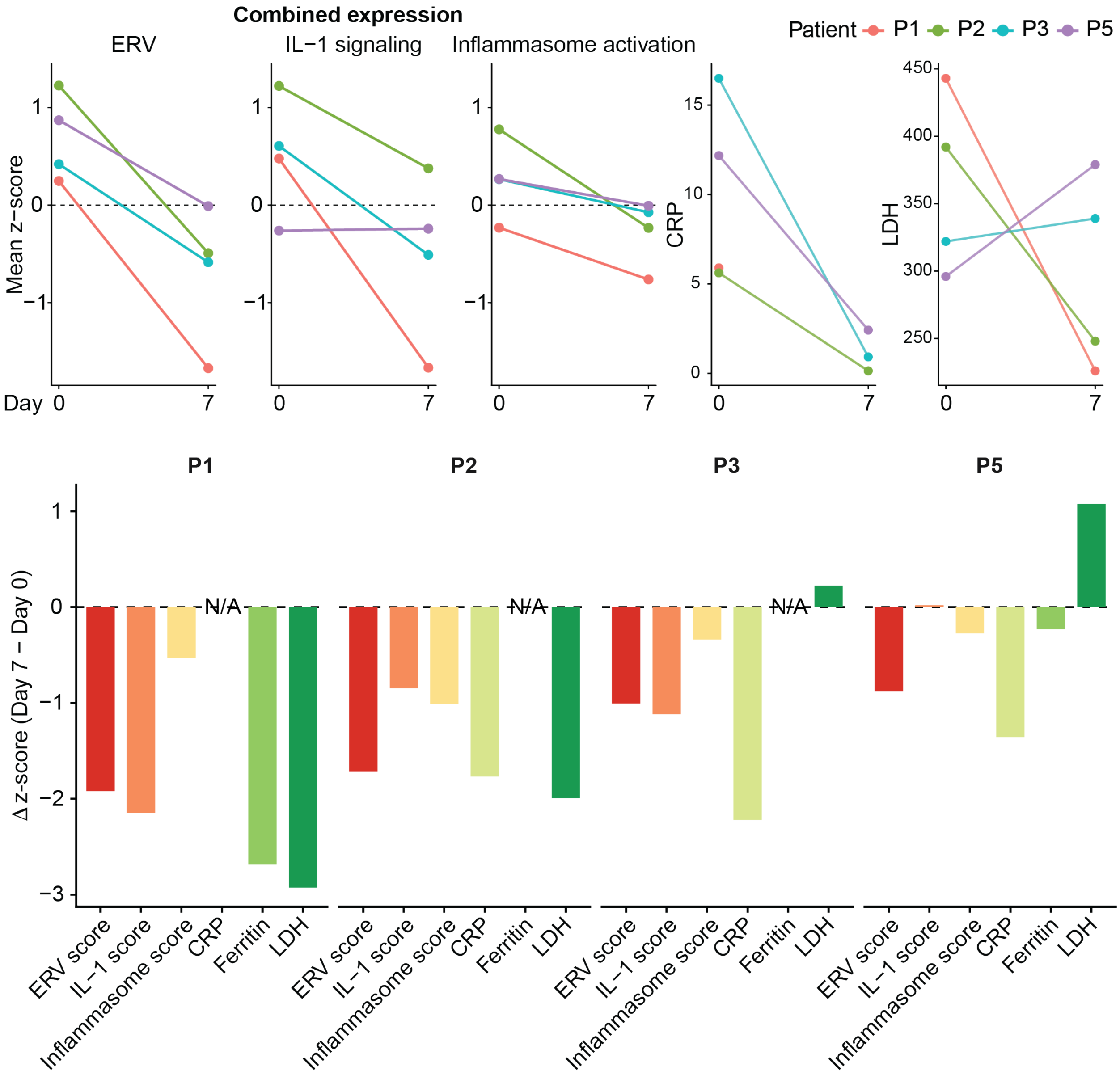


**Fig. S4. Decreases in ERV expression paralleled changes in inflammatory processes and clinical markers. Top:** Paired comparison of measurements at day 0 and day 7 for, from left to right, the combined expression of decreased signature ERVs, selected IL-1 signaling genes, NLRP3 inflammasome-associated genes, and the clinical markers CRP and LDH. **Bottom:** Changes in these features across individual patients, calculated as day 7 minus day 0 delta values.

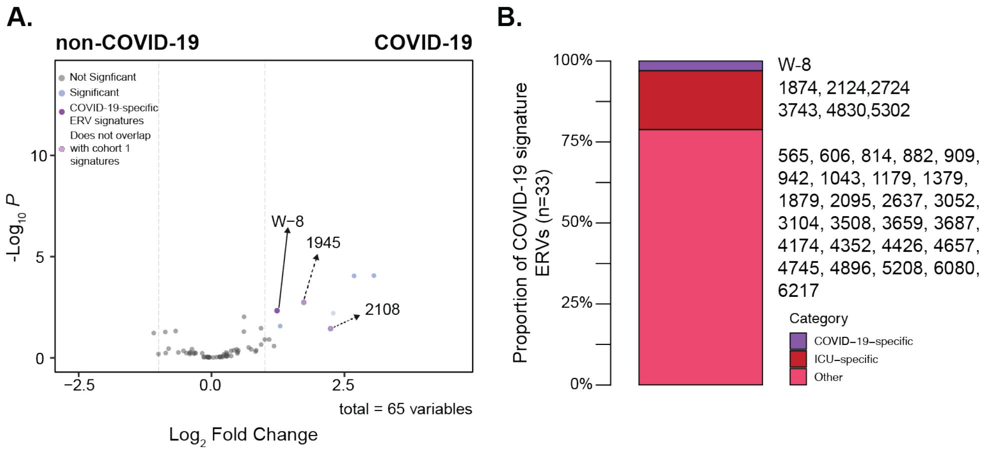


**Fig. S5. Identification of COVID-19-specific signature ERV. (A)** Volcano plot showing differential expression of ERVs between COVID-19 and non-COVID-19 patients. **(B)** Bar plot showing the proportion of severe COVID-19 signature ERVs that are ICU-specific and/or COVID-19-specific.

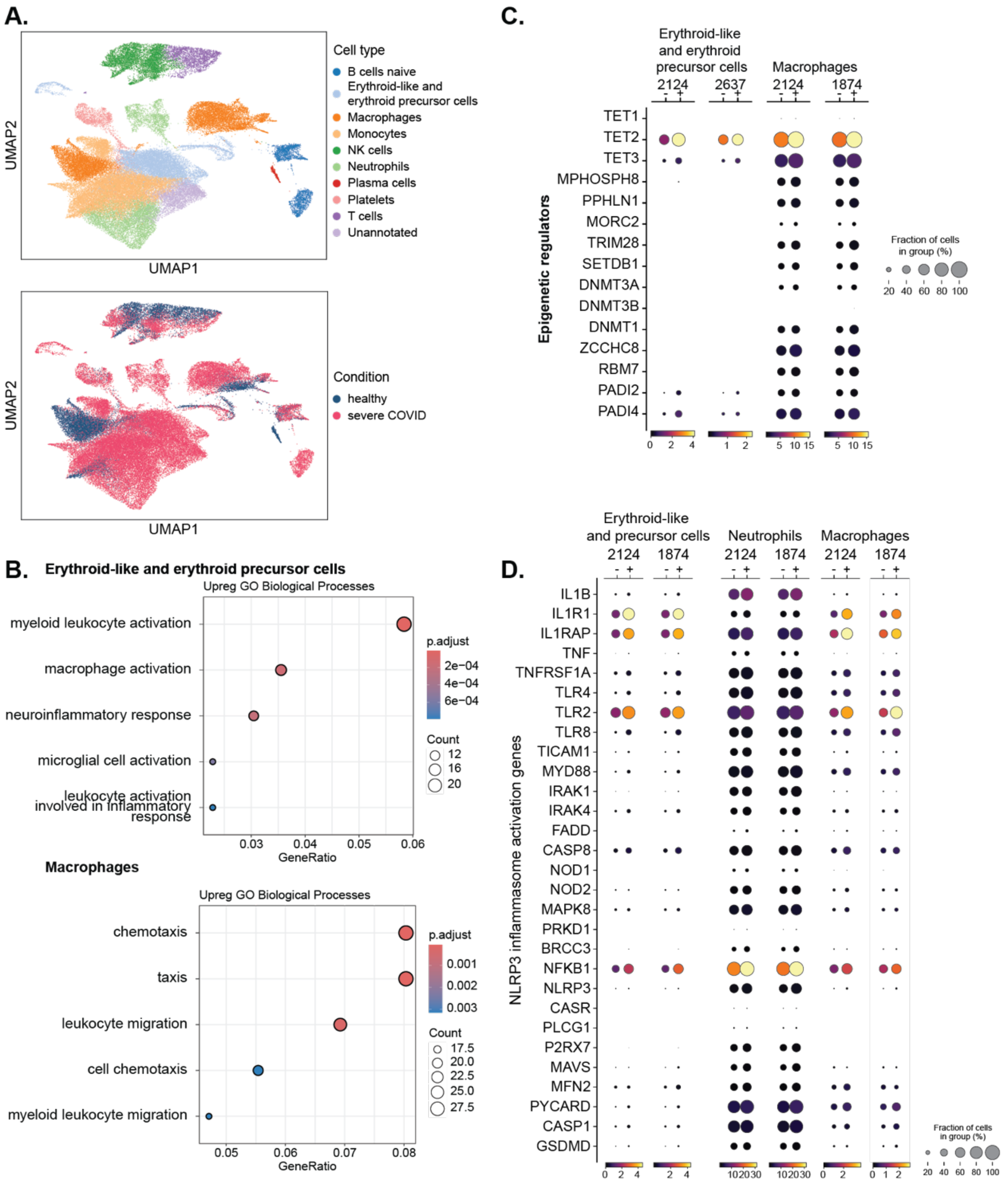


**Fig. S6. Immune cells expressing signature ERVs show heightened expression of innate immune activation genes. (A)** UMAP of whole-blood cells showing 14 clusters with cell type annotations (top) and disease conditions (bottom). **(B)** GO enrichment analysis of genes significantly upregulated in erythroid-like and erythroid precursor cells and macrophages from severe COVID-19 patients (adjusted p-value < 0.05, log2 fold change > 1). **(C)** Dot plots comparing the expression of epigenetic regulator genes in erythroid-like and erythroid precursor cells and macrophages that either express (+) or do not express (–) signature ERVs. (D) Dot plots comparing the expression of NLRP3 inflammasome activation genes in erythroid-like and erythroid precursor cells, neutrophils, and macrophages that either express (+) or do not express (–) signature ERVs.
